# BCHS acts as a stress transducer connecting autophagic quality control with DNA damage repair

**DOI:** 10.1101/2025.11.04.685471

**Authors:** Kuan-Hui Lu, Bo-Hua Yu, Tzu-Kang Sang

## Abstract

Cellular homeostasis relies on complex mechanisms that sense and respond to genotoxic stress. While autophagy adaptors are well-known for their role in removing cytoplasmic protein aggregates, their involvement in DNA damage responses is less clear. In this study, we show that Blue cheese (BCHS), the Drosophila homolog of mammalian Alfy/WDFY3, functions as a critical regulator connecting autophagy, DNA repair, and intercellular communication. Using a *Drosophila* eye model with TER94 (VCP ortholog) deficiency, we observed that genotoxic stress specifically increases expression and nuclear buildup of ref(2)P (p62/SQSTM1 homolog). Reducing BCHS worsens nuclear ref(2)P accumulation and DNA damage, while increasing BCHS levels alleviates these effects, highlighting BCHS as a vital regulator of nuclear proteostasis and genome stability. Mechanistically, genotoxic stress triggers reversible separation of the ref(2)P–BCHS complex, temporally coordinated with ref(2)P nuclear translocation. Strikingly, BCHS undergoes dynamic redistribution from the nuclear periphery to the extracellular matrix (ECM) via an exosome-dependent pathway involving Rab11, Rab27, Rab35, VAMP7, and tetraspanin Tsp96F. Blocking BCHS secretion significantly hampers DNA repair, demonstrating that extracellular BCHS export is functionally required for genome maintenance. These findings establish a paradigm wherein autophagy adaptors serve dual roles—coordinating nuclear stress responses while communicating cellular damage status to the tissue microenvironment. Given the conservation between BCHS and human Alfy, this mechanism may have broad implications for diseases characterized by defective proteostasis and genome instability, including neurodegeneration and cancer.

## Introduction

Cellular homeostasis relies on sophisticated surveillance mechanisms that identify and eliminate potentially harmful components. Autophagy, a highly conserved catabolic pathway in eukaryotic cells, is one such mechanism that sequesters cellular components inside double-membraned autophagosomes for degradation in lysosomes (or vacuoles in yeast), enabling the recycling of molecular building blocks during cellular stress^1^. The precise execution of autophagy depends on a coordinated network of autophagy-related genes (ATG) products that orchestrate these complex molecular processes^2^. Within this framework, selective autophagy operates as a specialized mechanism that employs core ATG alongside additional regulatory factors to identify and eliminate specific cellular components, including damaged organelles and protein aggregates, thereby preventing cellular dysfunction^3^. Central to this process is the targeted recruitment of cargo to the forming autophagosomes, mediated by autophagy receptors or adaptors that serve as molecular links between specific substrates and the autophagosomal machinery^4^.

Among these essential adaptors, Alfy (WDFY3) plays a critical role in aggrephagy, a selective autophagy pathway dedicated to removing protein aggregates^5,6^. Alfy orchestrates aggregate recognition and transfer to autophagosomes through its multi-domain structure. The C-terminal PH-BEACH domain mediates interaction with p62 (SQSTM1), a key autophagy receptor that recognizes ubiquitinated protein aggregates. Meanwhile, WD40 repeats facilitate direct interaction with ATG5, promoting recruitment of additional autophagosomal proteins (ATG12, ATG16, LC3) and linking cargo-bound p62 to membrane-associated LC3. The C-terminal FYVE domain, a specialized zinc finger motif, confers specificity for phosphatidylinositol 3-phosphate (PI(3)P), a lipid signaling molecule enriched on autophagic and endosomal membranes^5,7,8^. Notably, Alfy has been detected in the nucleus^9,10^, suggesting additional functions in processing nuclear components.

Studies of the *Drosophila* homolog Blue cheese (BCHS) have revealed conserved mechanisms underlying Alfy function. BCHS promotes efficient aggregate clearance through coordinated interactions with ATG5 and p62; its loss results in the pathological accumulation of ubiquitinated inclusions in neural tissues, leading to neurodegeneration^7,10^. These findings highlight the crucial role of this autophagy adaptor in maintaining cellular homeostasis and preventing pathological aggregation.

Our previous work revealed the multifaceted roles of Valosin-containing protein (VCP, known as TER94 in *Drosophila*), a conserved AAA+ ATPase that functions in both autophagy and DNA damage repair^11^. Loss of functional TER94 produced a distinctive nuclear expansion phenotype characterized by the accumulation of ubiquitinated proteins together with autophagy markers ATG8a and ref(2)P (*Drosophila* homolog of mammalian p62/SQSTM1) within the enlarged nuclear compartment. We also observed elevated γH2AV signals (*Drosophila* homolog of mammalian γH2AX), indicating accumulated DNA damage and compromised genomic integrity. These findings demonstrate that TER94 is essential for nuclear protein clearance and autophagy promotion while maintaining genomic stability, revealing complex interconnections between autophagy and DNA repair pathways.

Given Alfy’s nuclear localization and function as a p62 adaptor^12^, we investigated whether BCHS influences the nuclear phenotypes observed in TER94-deficient cells. Our study demonstrates that *Drosophila* BCHS functions as a critical sensor for genotoxic stress and reveals a novel mechanism linking autophagy regulation, DNA damage signaling, and intercellular communication. Following DNA damage, BCHS translocates from the nucleus to the extracellular space, triggering nuclear import of ref(2)P concomitant with elevated γH2AV levels. Our results establish that nuclear BCHS serves as a guardian of genomic integrity, with its exosome-mediated export to the extracellular space representing a previously unrecognized cellular response to genotoxic stress. This study provides new insights into how autophagy adaptors function beyond protein aggregate clearance, acting as integral components of the DNA damage response and revealing unexpected mechanistic connections between these fundamental cellular processes.

## Results

### TER94 deficiency triggers temporal changes in autophagy gene expression with sustained upregulation of ref(2)P

Our previous studies established that TER94 loss-of-function (LOF) cells exhibit elevated genotoxic stress and compromised autophagy. To investigate the molecular mechanisms underlying autophagy dysregulation in TER94-deficient cells, we utilized a *Drosophila* eye model expressing a dominant-negative TER94 mutant lacking ATP-binding capacity (*Rh1>TER94^K2A^*). We performed temporal expression profiling of autophagy-related genes at two critical developmental stages: 1-day-old flies, representing the early pre-symptomatic phase, and 5-day-old flies, when TER94 deficiency-induced nuclear expansion is morphologically evident^11^. This experimental design allowed us to distinguish early compensatory responses from sustained molecular alterations associated with cellular pathology.

Quantitative analysis revealed that at day 1, all examined autophagy genes displayed modest but consistent upregulation, with transcript levels increasing less than 2-fold relative to controls (Figure 1A and 1B). This coordinated induction is consistent with early activation of the autophagic machinery in response to TER94 impairment. Strikingly, by day 5, when nuclear morphology defects were prominent, expression of most autophagy genes had returned to near-baseline levels, suggesting transient activation followed by adaptation or exhaustion of the autophagic response (Figure 1A and 1C).

**Figure 1.**
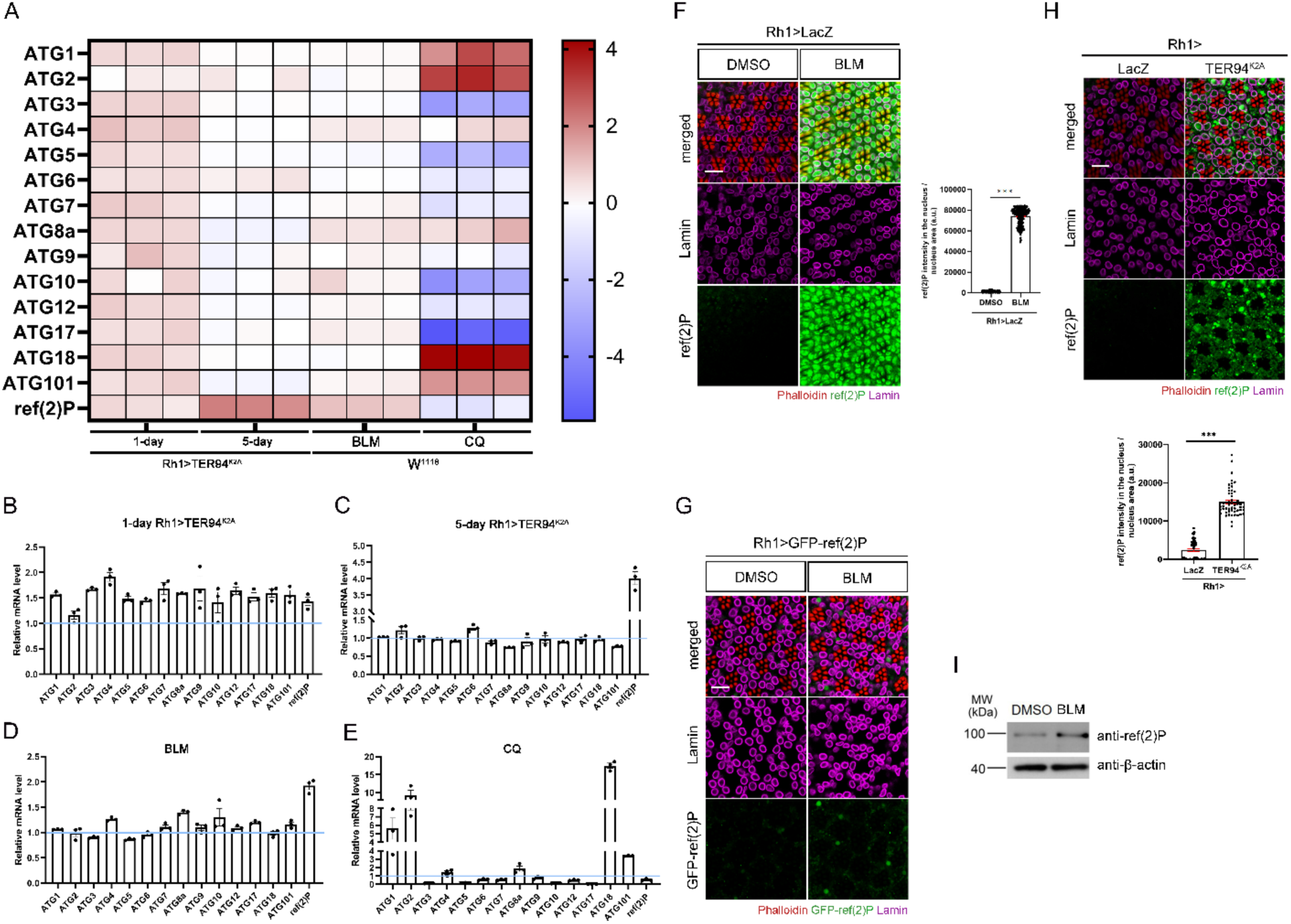
ref(2)P expression is upregulated and translocates to the nucleus in response to genotoxic stress. (A) Heatmap showing the relative mRNA expression (log₂ fold change) of indicated genes in 1-day-old and 5-day-old *Rh1>TER94^K2A^* flies, and *w^1118^* flies treated with bleomycin (BLM) or chloroquine (CQ). Color intensity represents the magnitude and direction of gene expression changes relative to control (blue, downregulation; red, upregulation). (B-E) Quantification of gene expression levels for groups shown in the heatmap. Data represent mean ± SEM from three biological replicates. (F) Left: Representative confocal images of 5-day-old *Rh1>LacZ* retinas treated with indicated drugs and immunostained for ref(2)P (green), Lamin (magenta), and phalloidin (red, F-actin). Right: Quantification of nuclear ref(2)P fluorescence intensity. (G) Representative confocal images of 5-day-old *Rh1>LacZ* retinas expressing GFP-ref(2)P treated with indicated drugs and stained with phalloidin (red) and anti-Lamin (magenta). (H) Top: Representative confocal images of 5-day-old control (*Rh1 > LacZ*) and *Rh1 > TER94^K2A^* retinas immunostained for ref(2)P (green), Lamin (magenta), and phalloidin (red). Bottom: Quantification of nuclear ref(2)P fluorescence intensity. (I) Western blot analysis of ref(2)P protein levels in *w^1118^* flies treated with indicated drugs. β-actin serves as a loading control. Scale bars, 10 µm.

In marked contrast, ref(2)P, the *Drosophila* ortholog of mammalian p62/SQSTM1, exhibited sustained and amplified upregulation, with transcript levels increasing approximately 4-fold at day 5 (Figure 1C). Unlike other autophagy components, ref(2)P expression remained elevated throughout disorder progression, indicating a persistent cellular stress response. These findings suggest that ref(2)P plays a critical and potentially unique role in mediating the cellular response to TER94 dysfunction, potentially through selective autophagy pathways that address the accumulation of ubiquitinated proteins in the nucleus or mitigate genotoxic stress, as previously demonstrated^11^.

### Genotoxic stress, rather than autophagy impairment, drives ref(2)P upregulation under TER94 deficiency

The sustained elevation of ref(2)P expression in TER94-deficient cells raised the question of which cellular stress predominantly drives this response. Given that TER94 LOF induces both genotoxic and proteotoxic stress, as evidenced by elevated nuclear levels of γH2AV and FK2, we sought to determine whether genotoxicity or autophagy dysfunction primarily drives ref(2)P induction. To dissect these contributions, we subjected wild-type (*w^1118^*) flies to either the DNA-damaging chemotherapeutic agent bleomycin (BLM, 70 µM) or the autophagy flux inhibitor chloroquine (CQ, 2.4 mM) for 24 hours prior to gene expression analysis.

BLM treatment recapitulated the ref(2)P response observed in TER94-deficient flies, eliciting approximately 2-fold upregulation of ref(2)P transcript levels, while other autophagy-related genes exhibited only minimal changes (Figure 1A and 1D). This selective induction pattern closely mirrored that observed in aged TER94^K2A^ flies, suggesting genotoxic stress as a primary trigger. In contrast, CQ treatment induced robust upregulation of core autophagy initiation machinery components—including ATG1, ATG2, ATG18, and ATG101—which function in early autophagosome formation (Figure 1A and 1E). Notably, autophagy inhibition failed to induce ref(2)P expression, despite effectively blocking autophagic flux. These pharmacological interventions demonstrate that ref(2)P induction is selectively responsive to genotoxic stress rather than generalized autophagy impairment in our model system.

### ref(2)P accumulates in nuclei following genotoxic stress

To validate these findings at the protein level, we examined both endogenous ref(2)P and an ectopically expressed GFP-ref(2)P reporter in *Rh1>TER94^K2A^* and BLM-treated flies. Immunofluorescence analysis revealed substantial increases in both endogenous ref(2)P and GFP-ref(2)P reporter signal following genotoxic insults (Figure 1F and 1G). Strikingly, under TER94 LOF conditions, a substantial fraction of ref(2)P protein exhibited nuclear localization (Figure 1H), a distribution pattern not typically observed under basal conditions. Western blot analysis corroborated these observations, demonstrating that BLM treatment elevated ref(2)P protein levels in wild-type flies (Figure 1I). Given that GFP-ref(2)P is degraded via autophagy, the accumulation of this reporter—together with converging evidence from transcriptional, immunofluorescence, and biochemical analyses—indicates that genotoxic stress simultaneously stimulates ref(2)P expression and impairs autophagic clearance, resulting in substantial protein accumulation.

### BCHS orchestrates ref(2)P nuclear translocation in response to genotoxic stress

Mammalian p62 cooperates with Alfy to mediate the clearance of protein aggregates through selective autophagy. The pronounced nuclear accumulation of ref(2)P and ubiquitinated proteins under TER94 LOF and genotoxic stress conditions led us to hypothesize that selective autophagy may be compromised in these conditions, and that BCHS—the *Drosophila* Alfy ortholog—might play a regulatory role in this process. To test this hypothesis, we depleted BCHS via RNAi in *Rh1>TER94^K2A^* flies. Significantly, BCHS depletion increased nuclear ref(2)P in both unstressed and TER94-deficient cells, demonstrating that BCHS actively prevents ref(2)P nuclear entry under both basal and stress conditions (Figure 2A). Furthermore, BCHS depletion in TER94-deficient photoreceptors enhanced nuclear accumulation of FK2-positive ubiquitinated proteins (Figure S1), directly linking nuclear ref(2)P buildup to impaired clearance of ubiquitinated nuclear substrates. Notably, BCHS reduction also elevated γH2AV levels (Figure 2B), suggesting that BCHS-mediated regulation of ref(2)P is functionally coupled to DNA damage resolution.

**Figure 2.**
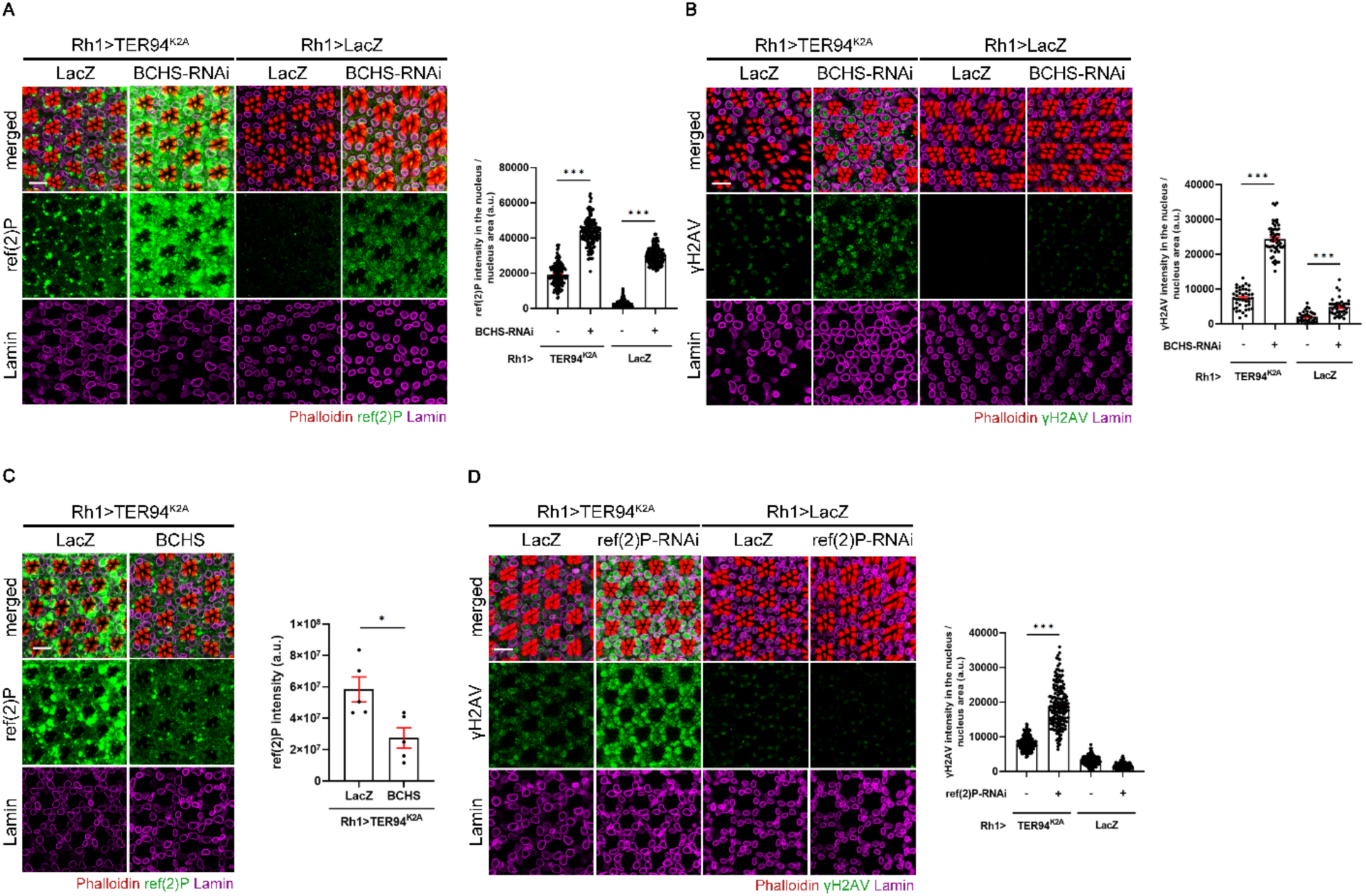
BCHS regulates the nuclear translocation of ref(2)P under genotoxic stress. (A) Left: Representative confocal images of 5-day-old control (*Rh1>LacZ*) and *Rh1>TER94^K2A^*retinas co-expressing *LacZ* or *BCHS-RNAi* immunostained for ref(2)P (green), Lamin (magenta), and phalloidin (red, F-actin). Right: Quantification of nuclear ref(2)P fluorescence intensity. (B) Left: Representative confocal images of 5-day-old control (*Rh1>LacZ*) and *Rh1>TER94^K2A^*retinas co-expressing *LacZ* or *BCHS-RNAi* immunostained for γH2AV (green), Lamin (magenta), and phalloidin (red). Right: Quantification of nuclear γH2AV fluorescence intensity. (C) Left: Representative confocal images of 5-day-old *Rh1>TER94^K2A^* retinas co-expressing *LacZ* or wild-type *BCHS* immunostained for ref(2)P (green), Lamin (magenta), and phalloidin (red). Right: Quantification of nuclear ref(2)P fluorescence intensity. (D) Left: Representative confocal images of 5-day-old control (*Rh1>LacZ*) and *Rh1>TER94^K2A^* retinas co-expressing *LacZ* or *ref(2)P-RNAi* immunostained for γH2AV (green), Lamin (magenta), and phalloidin (red). Right: Quantification of nuclear γH2AV fluorescence intensity. Scale bars, 10 μm.

Conversely, BCHS overexpression in *Rh1>TER94^K2A^* flies significantly reduced ref(2)P protein levels (Figure 2C), supporting a role for BCHS in promoting ref(2)P clearance or redistribution under stress. The functional importance of this regulatory axis was underscored by the observation that ref(2)P depletion exacerbated γH2AV accumulation in stressed cells (Figure 2D), suggesting that appropriate ref(2)P levels— potentially modulated by BCHS—are necessary for efficient DNA repair. Collectively, these genetic manipulations establish BCHS as a critical regulator of ref(2)P dynamics and genome stability under genotoxic stress conditions.

### Context-dependent dissociation of ref(2)P and BCHS during genotoxic stress

While mammalian Alfy functions as a scaffold adaptor that constitutively binds p62 to promote autophagic clearance of protein aggregates, our genetic data suggested that the ref(2)P–BCHS interaction might be dynamically regulated in response to genotoxic stress. To test whether BCHS modulation of nuclear ref(2)P involves changes in their physical association, we performed co-immunoprecipitation (co-IP) assays under three conditions: unstressed control, acute genotoxic stress (BLM treatment), and post-stress recovery.

Under basal conditions, coIP from nuclear protein revealed that ref(2)P and BCHS form a complex. Strikingly, BLM-induced genotoxic stress caused marked dissociation of the ref(2)P–BCHS complex, temporally coinciding with nuclear enrichment of ref(2)P (Figure 3). This stress-induced dissociation was reversible: following a 24-hour recovery period, the ref(2)P–BCHS interaction was substantially restored to baseline levels (Figure 3). This dynamic, context-dependent remodeling of the ref(2)P–BCHS complex suggests a novel regulatory mechanism wherein genotoxic stress triggers transient complex dissociation. We propose that this dissociation liberates ref(2)P for nuclear translocation and DNA damage response functions, while simultaneously repositioning BCHS for alternative stress-responsive roles.

**Figure 3.**
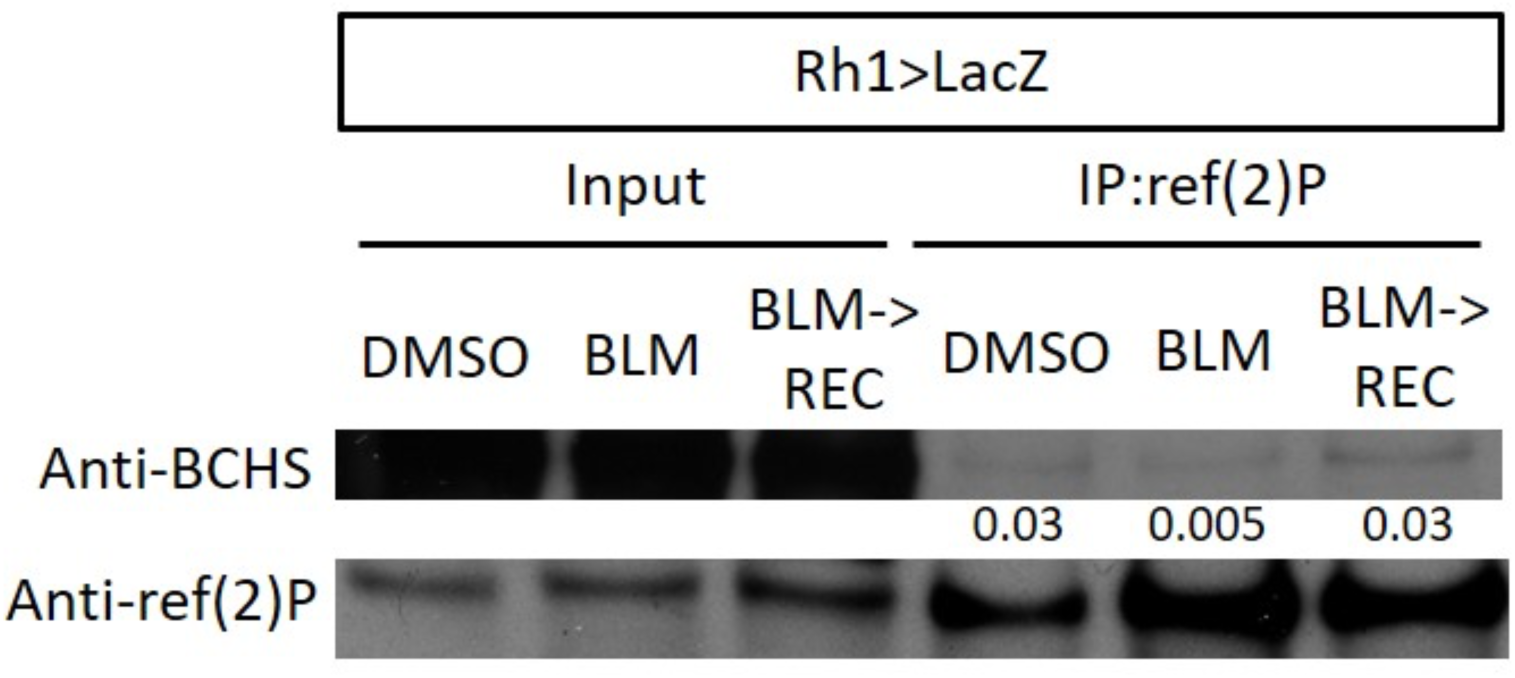
Context-dependent dissociation of ref(2)P and BCHS during genotoxic stress. Co-immunoprecipitation (co-IP) analysis showing the interaction between ref(2)P and BCHS under basal, genotoxic stress, and recovery conditions in *Rh1>LacZ* fly heads. Nuclear extracts were collected from flies treated with DMSO (control), bleomycin (BLM), or allowed to recover for 24 hours following bleomycin treatment (BLM → Rec). ref(2)P was immunoprecipitated from each sample, and both the input and IP fractions were immunoblotted with antibodies against BCHS and ref(2)P. The relative interaction level was quantified as the ratio of BCHS in the IP fraction to ref(2)P intensity in the same IP fraction.

### BCHS transitions from nuclear coordinator to extracellular cargo under genotoxic stress

Having established reversible dissociation of the ref(2)P–BCHS complex during genotoxic stress, we investigated how this separation influences BCHS subcellular localization. Both BCHS and mammalian Alfy have been reported to exhibit nuclear or perinuclear localization, prompting us to examine whether BCHS undergoes spatial reorganization under stress conditions. In unstressed *Rh1>LacZ* control photoreceptors, BCHS immunostaining revealed enrichment at the nuclear periphery (Figure 4A), consistent with a nuclear or perinuclear homeostatic function.

**Figure 4.**
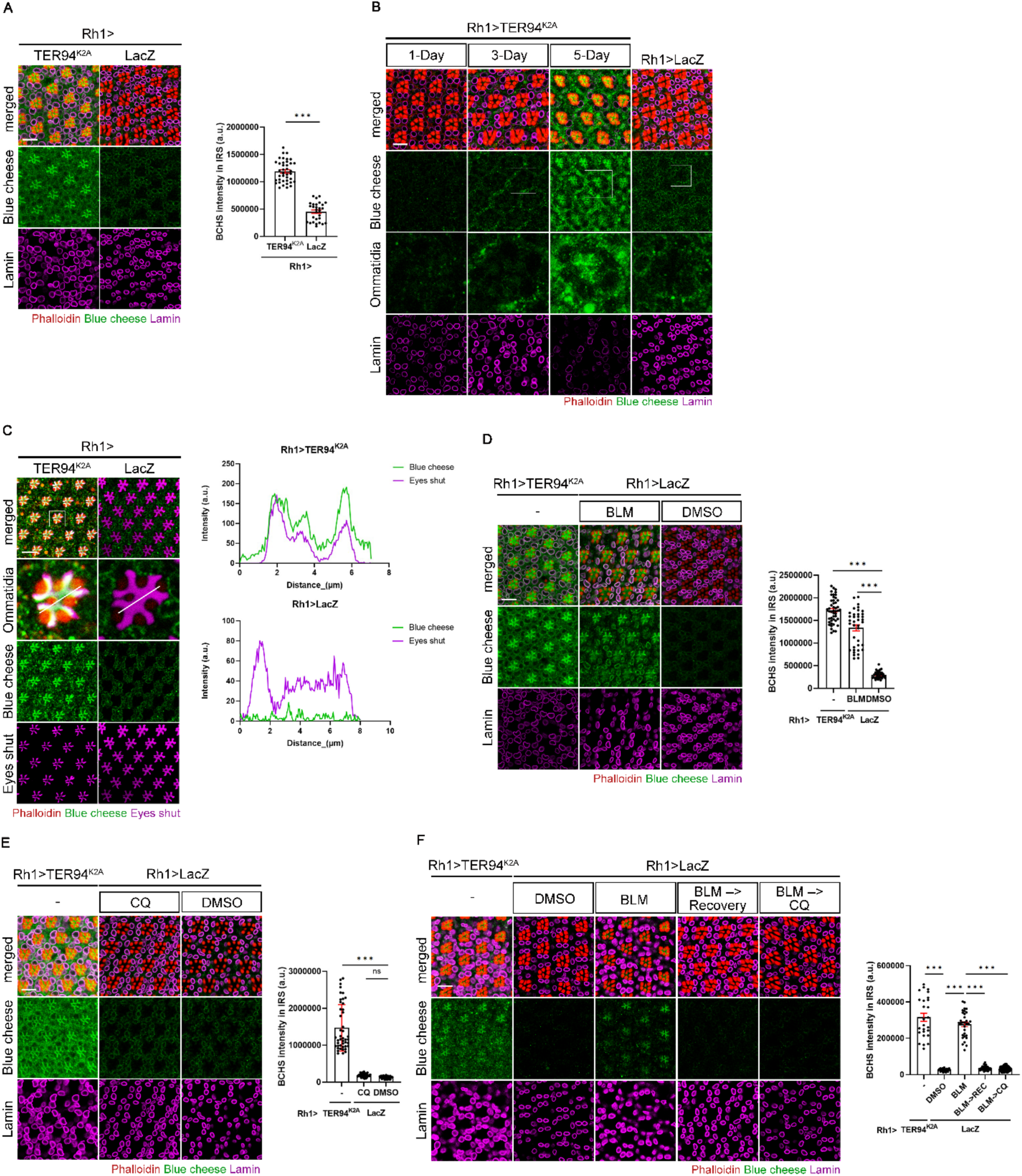
Genotoxic stress induces relocalization of BCHS from the nucleus to the extracellular space. (A) Left: Representative confocal images of 5-day-old *Rh1>LacZ* and *Rh1>TER94^K2A^*retinas immunostained for BCHS (green), Lamin (magenta), and phalloidin (red). Right: Quantification of BCHS fluorescence intensity in the interrhabdomeral space (IRS). (B) Time-course analysis of BCHS subcellular localization dynamics in *Rh1>TER94^K2A^* retinas at the indicated time points and a matched 5-day-old control (*Rh1>LacZ*). Retinas are immunostained for BCHS (green), Lamin (magenta), and phalloidin (red). Single ommatidia (white boxes) show BCHS staining only. (C) Left: Representative confocal images of 5-day-old *Rh1>LacZ* and *Rh1>TER94^K2A^* retinas immunostained for BCHS (green), Eys (magenta), and phalloidin (red). Single ommatidia from the indicated boxes are enlarged. Right: Fluorescence intensity line profiles of BCHS and Eys across the IRS. (D and E) Left: Representative confocal images of 5-day-old *Rh1>LacZ* retinas treated with indicated drugs or *Rh1>TER94^K2A^* retinas immunostained for BCHS (green), Lamin (magenta), and phalloidin (red). Right: Quantification of BCHS fluorescence intensity in the IRS. (F) Left: Representative confocal images of 5-day-old *Rh1>LacZ* control retinas treated with indicated drugs or *Rh1>TER94^K2A^* retinas. Control flies were treated with DMSO or BLM. One day without drug treatment following BLM exposure is labeled "BLM → Recovery", and one day of CQ feeding after BLM treatment is labeled "BLM → CQ." Retinas are immunostained for BCHS (green), Lamin (magenta), and phalloidin (red). Right: Quantification of BCHS fluorescence intensity in the IRS. Scale bars, 10 μm.

Remarkably, TER94-deficient *Rh1>TER94^K2A^* photoreceptors exhibited dramatic BCHS redistribution to extracellular compartments (Figure 4A). Time-course analysis of 1-day, 3-day, and 5-day-old *Rh1>TER94^K2A^* retinas revealed progressive BCHS relocalization from the nuclear periphery to both basolateral and apical extracellular matrix (ECM). By day 5, BCHS displayed a distinct pattern of accumulation within the interrhabdomeral space (IRS)–the specialized apical ECM between photoreceptor rhabdomeres—in each ommatidium (Figure 4B). Co-immunostaining with eyes shut (eys), an IRS-specific matrix protein, confirmed complete colocalization (Figure 4C), definitively establishing BCHS secretion into this extracellular compartment. These findings reveal a previously unrecognized stress-induced subcellular redistribution of BCHS from nuclear-associated functions to an extracellular destination, suggesting a novel non-cell-autonomous role for this autophagy regulator.

To determine whether genotoxic stress specifically triggers BCHS redistribution, we treated *Rh1>LacZ* control flies with BLM for 24 hrs to induce DNA damage. While vehicle (DMSO)-treated controls displayed normal nuclear/perinuclear BCHS localization, BLM treatment induced pronounced protein concentration in the IRS (Figure 4D). Importantly, this redistribution appeared genotoxicity-specific rather than a general stress response, as CQ-mediated autophagy inhibition did not provoke BCHS secretion (Figure 4E), despite effectively blocking autophagy flux.

We next examined whether secreted BCHS persisted in the IRS following resolution of genotoxic stress. Strikingly, when BLM-treated flies were allowed a 24-hour recovery, BCHS signals in the IRS were no longer detectable (Figure 4F), mirroring the temporal dynamics of ref(2)P–BCHS complex reassociation observed in our co-IP experiments (Figure 3). This clearance of BCHS from the IRS occurred independently of autophagy, as treatment with CQ during the recovery phase did not prevent BCHS removal (Figure 4F). Together, these findings demonstrate that genotoxicity triggers a transient, reversible relocation of BCHS to extracellular compartments, representing a dynamic stress-responsive mechanism that is spatially and temporally coordinated with ref(2)P–BCHS complex dissociation and nuclear ref(2)P accumulation.

### The exosome pathway mediates genotoxicity-induced BCHS secretion

BCHS lacks a typical signal peptide for conventional secretion, suggesting that vesicle-mediated transport likely facilitates its export from the cell. To test this hypothesis, we systematically knocked down various Rab GTPases and found that suppression of Rab11, Rab27, or Rab35— but not Rab3, Rab5, Rab6, Rab7, and Rab14—prevented BCHS from moving to the IRS in stressed *Rh1>TER94^K2A^*flies (Figure 5A-C, and Figure S2). Notably, Rab11, Rab27, and Rab35 are all associated with exosome secretion^13^, prompting us to examine additional components of this pathway. We then depleted VAMP7 (vesicle-associated membrane protein 7), a key R-SNARE involved in the fusion of multivesicular bodies (MVBs) with the plasma membrane^14^. Consistent with exosome-dependent secretion, VAMP7 knockdown blocked BCHS accumulation in the IRS (Figure 5D).

**Figure 5.**
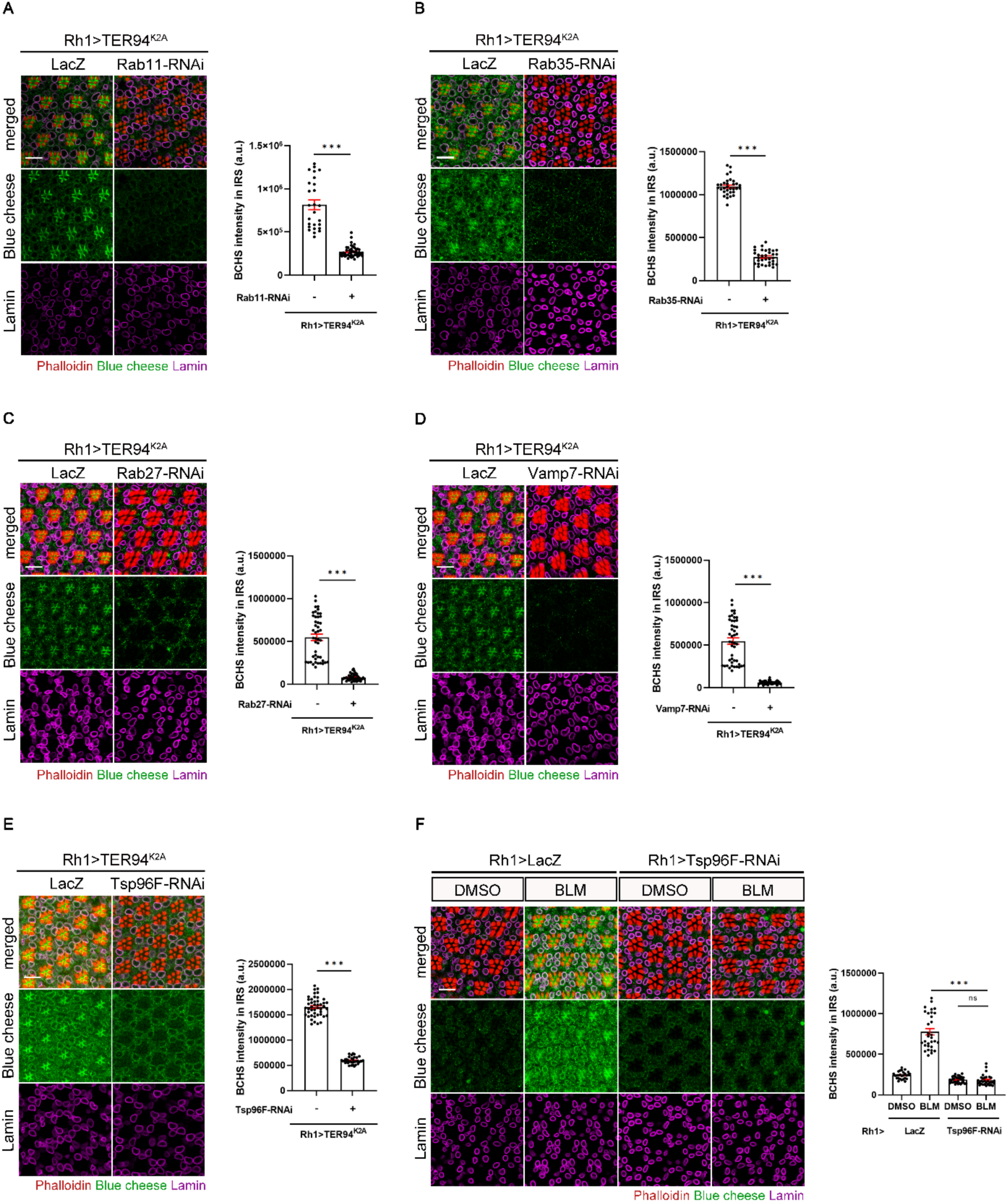
Genotoxic stress promotes BCHS relocalization to the extracellular matrix via the exosome pathway. (A-E) Left: Representative confocal images of 5-day-old *Rh1>TER94^K2A^*retinas co-expressing *LacZ* or indicated knockdown constructs immunostained for BCHS (green), Lamin (magenta), and phalloidin (red). Right: Quantification of BCHS fluorescence intensity in the IRS. (F) Left: Representative confocal images of 5-day-old *Rh1>LacZ* and *Rh1>Tsp96F-RNAi* retinas treated with DMSO or BLM immunostained for BCHS (green), Lamin (magenta), and phalloidin (red). Right: Quantification of BCHS fluorescence intensity in the IRS. Scale bars, 10 μm.

To further validate exosome-dependent BCHS secretion, we knocked down Tsp96F, a *Drosophila* tetraspanin homolog essential for exosome biogenesis and cargo sorting. Tetraspanins organize microdomains within MVBs and facilitate their fusion with the plasma membrane. Consistent with this function, Tsp96F knockdown in *Rh1>TER94^K2A^* flies abolished BCHS secretion to the IRS while preserving intracellular BCHS levels (Figure 5E). Similarly, BLM treatment of *Rh1>Tsp96F-RNAi* flies failed to induce BCHS accumulation in the IRS, despite maintaining intracellular BCHS signals (Figure 5F). These results confirm that Tsp96F loss specifically blocks the secretory process without affecting BCHS expression or stability.

Together with the requirement for Rab11, Rab27, Rab35, and VAMP7, these findings demonstrate that the exosome pathway mediates the secretion of BCHS in response to genotoxic stress.

### Preventing BCHS secretion to the ECM impairs DNA damage repair

Accumulating evidence suggests that ECM mechanical properties can influence DNA damage repair efficiency. Given that BCHS is secreted to the ECM under genotoxic stress, we investigated whether this redistribution affects genome stability. We compared γH2AV levels in BLM-treated flies that either permit (*Rh1>LacZ)* or block (*Rh1>Tsp96F-RNAi)* BCHS secretion, examining retinas immediately after treatment and following 24 hrs of recovery.

In *Rh1>LacZ* control retinas, γH2AV signals increased immediately post-treatment but declined substantially after recovery (Figure 6), coinciding with clearance of BCHS from the ECM (Figure 4D). Strikingly, *Rh1>Tsp96F-RNAi* retinas exhibited significantly elevated γH2AV levels after the 24-hour recovery period compared to immediately post-treatment (Figure 6). Since Tsp96F knockdown prevents BCHS secretion (Figure 5F), these findings indicate that blocking BCHS export to the ECM impairs DNA damage repair.

**Figure 6.**
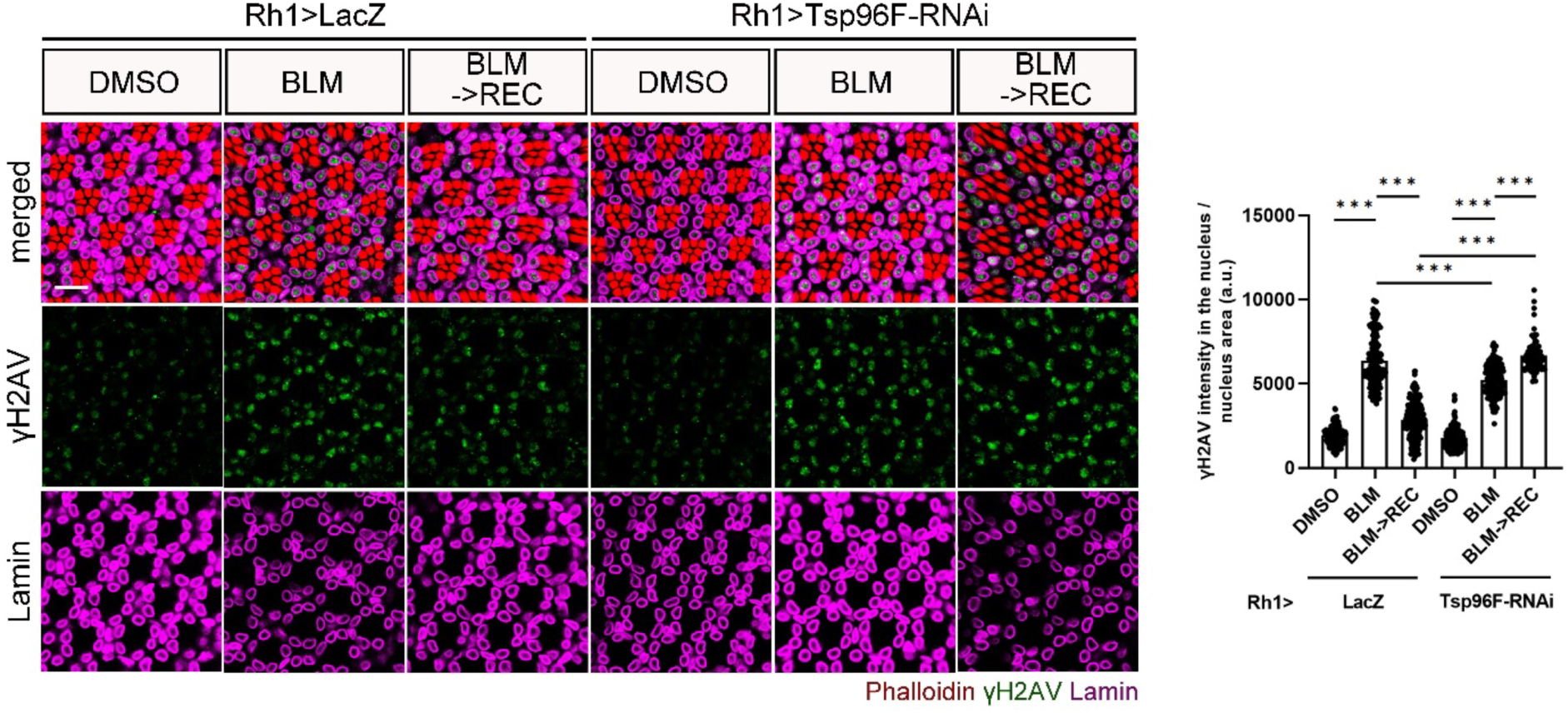
Inhibition of BCHS secretion into the ECM impairs the repair of DNA damage. Left: Representative confocal images of 5-day-old *Rh >LacZ* (control) and *Rh1>Tsp96F-RNAi* retinas under three conditions: DMSO treatment (vehicle control), BLM treatment (bleomycin-induced DNA damage), or one day of recovery on normal food following BLM treatment (BLM → Rec). Retinas were immunostained for BCHS (green), Lamin (magenta), and phalloidin (red). Right: Quantification of γH2AV fluorescence intensity in the nuclei. Scale bar, 10 µm

## Discussion

Using the *Drosophila* eye model, we have investigated the functional importance of VCP in maintaining genome stability and autophagy operation^11^. Here, we identify *Blue cheese (BCHS)* as a critical molecular link connecting DNA damage responses and autophagy. We demonstrate that ref(2)P, the *Drosophila* homolog of p62/SQSTM1, undergoes transcriptional induction and nuclear relocalization in response to TER94 (Drosophila VCP ortholog) dysfunction or direct genotoxic insult. Critically, our data identify BCHS as a crucial regulator of this nuclear ref(2)P dynamics. BCHS depletion exacerbated ref(2)P and ubiquitin accumulation within the nucleus and heightened γH2AV levels under both TER94-deficient and genotoxic conditions, whereas BCHS overexpression ameliorated these phenotypes. These findings position BCHS upstream of ref(2)P, where it coordinates nuclear protein quality control with DNA damage repair.

The nuclear enrichment of ref(2)P under genotoxic stress reveals that the nucleus is not merely a passive target of DNA damage but an active site where selective autophagy components participate in stress mitigation. This nuclear involvement aligns with emerging evidence that autophagic factors operate outside canonical cytoplasmic contexts to regulate chromatin integrity^15^, nuclear protein turnover^16,17^, and repair of DNA lesions^18^. For instance, LC3 contributes to double-strand break (DSB) resolution by interacting with chromatin and promoting recruitment of repair proteins^19^, while p62 modulates homologous recombination by sequestering RNF168 and limiting chromatin ubiquitination^18^. Although autophagy adaptors such as Alfy/WDFY3 and p62 are best characterized for their cytoplasmic roles in aggregate clearance^6,12^, several studies have hinted at their nuclear presence and possible signaling functions^9,10^. Our work expands this conceptual framework by establishing BCHS as a nuclear-localized autophagy adaptor that directly participates in genotoxic stress responses.

Beyond its nuclear role, our findings reveal that BCHS functions as a dynamic regulator that translocates between nuclear and extracellular compartments in response to genotoxic stress. Under basal conditions, BCHS associates with the nuclear periphery, where it may surveil nuclear proteostasis. Upon DNA damage, BCHS undergoes relocalization to the extracellular matrix (ECM) through an exosome-dependent mechanism involving Rab11, Rab27, Rab35, VAMP7, and the tetraspanin Tsp96F. This stress-induced translocation is not a passive event but a regulated process essential for efficient DNA damage repair. Preventing BCHS secretion through Tsp96F knockdown leads to sustained γH2AV accumulation, indicating defective DNA damage resolution and demonstrating that BCHS export is functionally required for genome maintenance. This observation establishes a previously unrecognized aspect of autophagy adaptor functionality. BCHS serves as a stress-responsive switch that connects genome stability with external signaling when faced with genotoxic stress.

This dynamic redistribution aligns with emerging evidence that components of the autophagy and endolysosomal machinery can be repurposed for unconventional secretion^20–22^. Notably, recent work demonstrated that DNA damage enhances secretion of autophagy-related proteins and small extracellular vesicles (sEVs) carrying cytoprotective or inflammatory signals^20,23^. Secreted autophagy proteins, including LC3 and ATG8 homologs, have been detected in extracellular vesicles where they influence immune modulation and stress adaptation^24,25^. BCHS could represent a parallel mechanism whereby autophagy adaptors communicate cellular stress status to the tissue microenvironment.

The requirement for BCHS export in DNA damage resolution raises important mechanistic questions. One possibility is that BCHS secretion relieves the nucleus from stress-induced protein burden, thereby facilitating recovery of DNA repair capacity. Our previous work established that loss of TER94 impairs both autophagy and DNA repair, resulting in nuclear accumulation of ubiquitinated proteins and autophagic markers. The current findings position BCHS as an effector downstream or parallel to TER94 that coordinates nuclear proteostasis with stress-induced communication. VCP/TER94 facilitates chromatin-associated degradation and DNA repair through extraction of ubiquitinated substrates from chromatin^26^. The BCHS–ref(2)P axis may act as a complementary route that handles persistent nuclear aggregates and signals through the exosome pathway when proteostasis cannot be restored intracellularly.

Alternatively, extracellular BCHS may serve a signaling function, modulating the tissue environment to support coordinated repair responses. This possibility is supported by accumulating evidence linking ECM properties to DNA repair efficiency. Mechanical cues and matrix integrity can modulate chromatin organization and repair through mechanotransduction^27,28^. By contributing to ECM remodeling under stress, BCHS secretion may indirectly influence nuclear responses, linking external and internal stress sensing. The correlation between BCHS relocalization to the ECM and nuclear import of ref(2)P further suggests coordinated regulation between these compartments during the genotoxic response.

Conceptually, this study redefines the autophagy adaptor paradigm by unveiling a dual nuclear–extracellular role for BCHS in genome stability maintenance. Mechanistically, it integrates nuclear autophagy, exosome biology, and DNA repair into a unified stress-response framework. Given the conservation between BCHS and human Alfy/WDFY3, this mechanism may have broader implications for diseases where both DNA damage and exosome secretion are dysregulated. In neurodegenerative disorders, impaired Alfy function leads to aggregate accumulation and neuronal loss^9,29^. In cancer, exosome-mediated stress signaling modulates therapy resistance and tumor progression. BCHS/Alfy-mediated export under DNA damage may thus represent a conserved adaptive strategy that balances intracellular repair with intercellular communication, maintaining tissue homeostasis under genotoxic stress. Understanding this pathway could reveal new therapeutic targets for conditions characterized by defective proteostasis and genome instability.

## Materials and Methods

### *Drosophila* genetics

Flies were raised on standard cornmeal food at 25°C under a 12-hour light/dark cycle. The following *Drosophila* strains were used: *Rh1-GAL4* (provided by Dr. Larry Zipursky), *UAS-LacZ*, *UAS-TER94^K2A^*, as previously described^30^. All transgenic RNAi lines were obtained from the Bloomington Drosophila Stock Center (Bloomington, IN, USA) or the Vienna Drosophila Resource Center (VDRC).

### Immunohistochemistry

Whole-mount preparations of fly eyes were performed as previously described^31^. The following primary antibodies were used: rabbit anti-BCHS (1:500, kindly provided by Dr. Rachel Kraut, Max-Planck-Institute of Molecular Cell Biology and Genetics), mouse anti-Lamin Dm0 (1:20, ADL67.10, Developmental Studies Hybridoma Bank, DSHB), rabbit anti-Lamin Dm0 (1:2000, generous gift from Dr. Paul Fisher), mouse anti-mono- and polyubiquitinated conjugates (FK2) (1:100, PW8810-0100, Enzo Life Sciences), rabbit anti-ref(2)P (1:20, ab178440, Abcam), and mouse anti-γH2Av (1:200, UNC93-5V2.1, DSHB).

Secondary antibodies were used at 1:100 dilution and included Alexa Fluor® 488, Alexa Fluor® 647, Cy3, and Cy5 conjugated secondary antibodies: Alexa Fluor® 488 AffiniPure Goat Anti-Mouse IgG (H+L) (115-545-146, Jackson ImmunoResearch Laboratories), Alexa Fluor® 488 AffiniPure Goat Anti-Rabbit IgG (H+L) (111-545-144, Jackson ImmunoResearch Laboratories), Alexa Fluor® 647 AffiniPure Goat Anti-Mouse IgG (H+L) (115-605-146, Jackson ImmunoResearch Laboratories), and Cy™5 AffiniPure Goat Anti-Rabbit IgG (H+L) (111-175-144, Jackson ImmunoResearch Laboratories). F-actin-enriched rhabdomeres were labeled with Rhodamine-conjugated phalloidin (1:20, P1951, Sigma-Aldrich).

Fluorescent images were acquired using a Zeiss LSM-510 or LSM-780 confocal microscope and processed with Adobe Photoshop. For comparative analyses among genotypes, all samples were prepared, imaged, and processed under identical conditions.

### Extraction of cytosolic and nuclear proteins from *Drosophila* retina

Cytosolic and nuclear protein fractions were sequentially extracted from *Drosophila* retina as previously described with minor modifications^11,32^. Briefly, freshly dissected or snap-frozen retinas (typically 100–200 pairs per sample) were collected in pre-chilled 1.5 mL microcentrifuge tubes on ice and homogenized in 100–200 μL of cold cytosolic extraction buffer (10 mM HEPES, pH 7.9; 10 mM KCl; 0.1 mM EDTA; 0.1 mM EGTA; 0.1% NP-40; 1 mM DTT; supplemented with protease and phosphatase inhibitors). Homogenates were incubated on ice for 10–15 min with gentle mixing and centrifuged at 1,500 × *g* for 5 min at 4°C. The supernatant was collected as the cytosolic fraction.

The remaining pellet containing nuclei was resuspended in 50–100 μL of cold nuclear extraction buffer (20 mM HEPES, pH 7.9; 0.4 M NaCl; 1 mM EDTA; 1 mM EGTA; 1% Triton X-100; 1 mM DTT; supplemented with protease and phosphatase inhibitors), vortexed for 15–30 s, and incubated on ice for 15–30 min with intermittent mixing. Samples were centrifuged at 14,000 × *g* for 10 min at 4°C, and the supernatant was collected as the nuclear fraction.

Protein concentrations were determined using the Bradford or BCA assay. Equal amounts of cytosolic and nuclear proteins were mixed with SDS sample buffer for SDS– PAGE and immunoblotting. All steps were performed on ice or at 4°C to minimize protein degradation.

### Co-immunoprecipitation

For co-immunoprecipitation, 100 fly heads were collected and homogenized in cold PBS containing protease inhibitors. Lysates were incubated with ref(2)P antibody at 4°C overnight, followed by incubation with protein A beads (Sigma-Aldrich) for 1 hr at room temperature. Beads were washed extensively, and bound proteins were eluted for SDS-PAGE and subsequent immunoblot analysis.

### Immunoblotting

For immunoblot analysis, the following primary antibodies were used: rabbit anti-BCHS (1:6000), rabbit anti-ref(2)P (1:2000, ab178440, Abcam), and anti-ß-Actin (1:20,000, GeneTex). HRP-conjugated secondary antibodies (GeneTex) were used at 1:10,000 dilutions. All loading controls were prepared by stripping the reagents from the original membrane and then re-immunoblotting the indicated antigen using standard procedures.

### BCHS quantification in the interrhabdomeral space (IRS)

Quantification of BCHS signal in the IRS was performed using ImageJ particle analysis. Regions of interest (ROIs) were defined by drawing circular areas corresponding to the distance between the R1 and R7 photoreceptor edges. BCHS fluorescence intensity within each ROI was measured and expressed in arbitrary units (a.u.).

### RT-qPCR

Total RNA was extracted from the eyes of *Rh1 > LacZ*, *Rh1 > TER94^K2A^*, and drug-treated flies using the GENEzol™ TriRNA Pure Kit (Geneaid) following the manufacturer’s instructions. RNA concentration and purity were determined spectrophotometrically. For cDNA synthesis, 5 μg of total RNA was reverse-transcribed using SuperScript II Reverse Transcriptase (Invitrogen) according to the manufacturer’s protocol. Quantitative PCR (qPCR) was performed using 1 μg of cDNA and gene-specific primers. Expression levels of autophagy-related genes were normalized to the internal control rp49. Primer sequences are provided in Supplementary Table 1. Relative expression levels were calculated using the 2^−ΔΔ*Ct*^ method.

### Drug administration

Drug treatments were administered by incorporating the specified compounds into standard fly food. Bleomycin (71 μM) and Chloroquine (2400 μM) were freshly prepared and thoroughly mixed into molten food cooled to 25°C before dispensing. To confirm ingestion, a trace amount of blue or red food dye was added to both drug-containing and control media, and food intake was verified by visualizing dye accumulation in the abdomens of treated flies. Adult flies were transferred onto the prepared media and maintained for 24 hrs at 25°C under standard light–dark conditions. Following treatment, flies were immediately collected for subsequent analyses.

### Statistical analysis

Data are presented as the mean ± standard error of the mean (SEM). Statistical significance between two groups was determined using Student’s *t*-test. Comparisons among more than two groups were analyzed using one-way ANOVA followed by Bonferroni’s post hoc test. Significance levels were denoted as *p<0.05* (*), *p<0.01* (**), and *p<0.001* (***).

## Acknowledgements

We thank Dr. Rachel Kraut for generously providing the anti-BCHS antibody. We also thank the Bloomington Drosophila Stock Center, Vienna *Drosophila* RNAi Center, and Fly Core Taiwan for supplying fly strains. Our gratitude extends to the Image Core of the Brain Research Center at National Tsing Hua University for assistance with confocal microscopy. We appreciate Dr. Henry Chang at Purdue University for helpful comments and discussions regarding this work. This research was supported by grants from the National Science and Technology Council in Taiwan (111-2311-B-007-007-MY3 and 114-2311-B-007-008).

## Author contributions

Kuan-Hui Lu and Tzu-Kang Sang conceived and designed the experiments. Kuan-Hui Lu, Bo-Hua Yu, and Tzu-Kang Sang conducted the experiments. Kuan-Hui Lu, Bo-Hua Yu, and Tzu-Kang Sang analyzed the data. Kuan-Hui Lu and Tzu-Kang Sang wrote the paper.

---

**Supplementary Figure 1. Nuclear accumulation of ref(2)P correlates with reduced degradation of ubiquitinated nuclear proteins.** Left: Representative confocal micrographs of 5-day-old control (*Rh1*>*LacZ*) and *Rh1*>*TER94*^K2A^ adult retinas co-expressing *LacZ* (control) or *BCHS-RNAi*. Retinas were immunostained for FK2 (ubiquitinated proteins, green), Lamin (magenta), and phalloidin (red). Right: Quantification of FK2 fluorescence intensity in the nuclei. Scale bar, 10 µm.

**Supplementary Figure 2. Rab GTPases associated with vesicle trafficking pathways other than exosome secretion do not affect BCHS secretion.** (A-E) Left: Representative confocal micrographs of 5-day-old *Rh1*>*TER94*^K2A^ adult retinas co-expressing *LacZ* (control) or the indicated *Rab-RNAi* constructs. Retinas were immunostained for BCHS (green), Lamin (magenta), and phalloidin (red). Right: Quantification of BCHS fluorescence intensity in the interrhabdomeral space (IRS). Scale bar, 10 µm.

