## Supplementary figures and images for "BCHS acts as a stress transducer connecting autophagic quality control with DNA damage repair"

### Supplemental Figures

Supplementary Figure 1

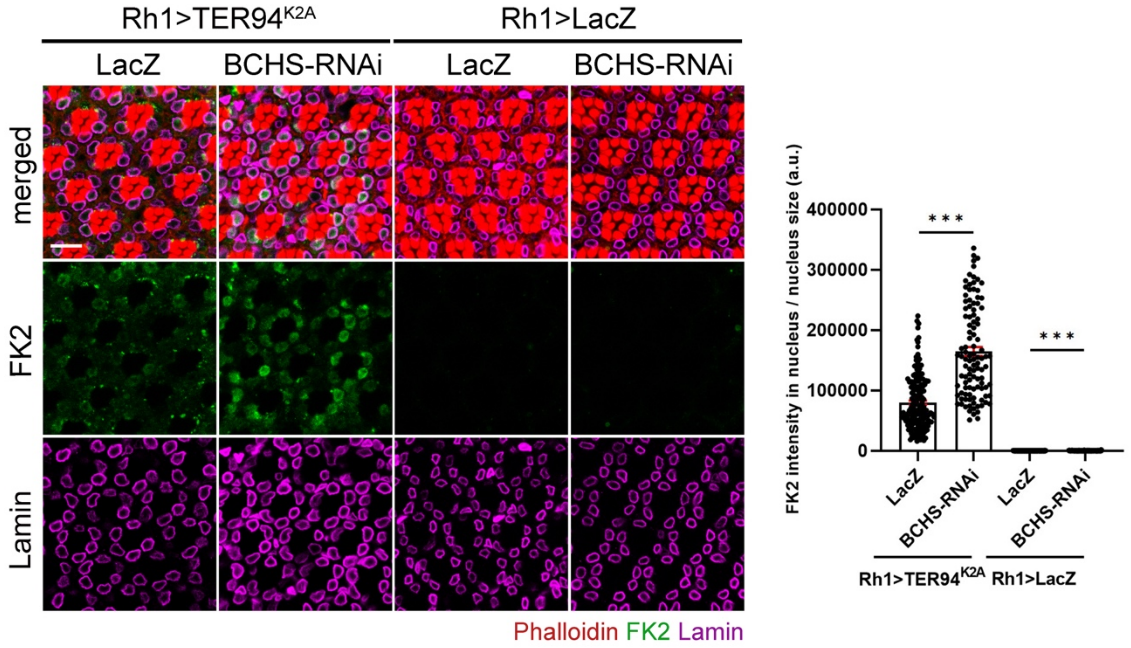

Supplementary Figure 2

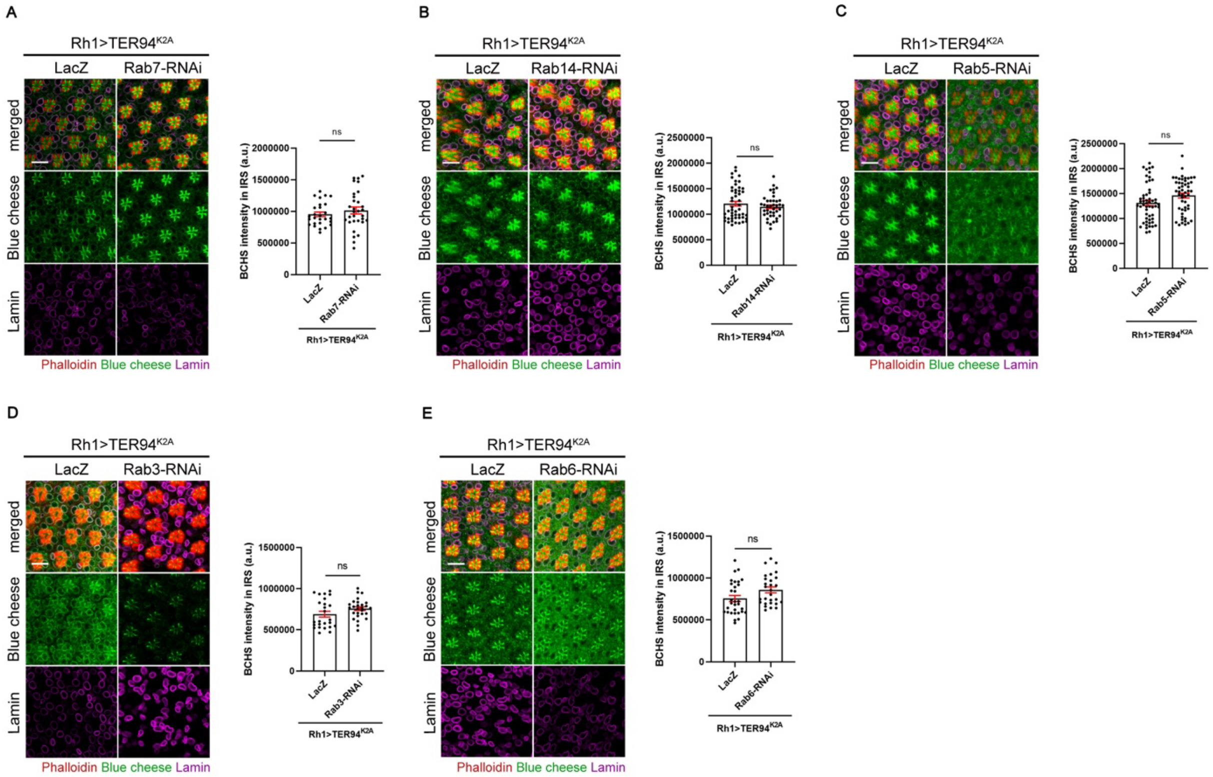
